# Neuroscientific Insights into the Built Environment: A Systematic Review of Empirical Research on Indoor Environmental Quality, Physiological Dynamics, and Psychological Well-Being in Real-Life Contexts

**DOI:** 10.1101/2025.03.11.642386

**Authors:** Aitana Grasso-Cladera, Maritza Arenas-Pérez, Paulina Wegertseder-Martinez, Erich Vilina, Josefina Mattoli-Sánchez, Francisco J. Parada

## Abstract

The research aims to systematize the current scientific evidence on methodologies used to investigate the impact of indoor built environment on well-being, focusing on Indoor Environmental Quality (IEQ) variables such as thermal comfort, air quality, noise, and lighting. This systematic review adheres to the Joanna Briggs Institute framework and PRISMA guidelines to assess empirical studies that incorporate physiological measurements like heart rate, skin temperature, and brain activity, captured through various techniques in real-life contexts.

The principal results reveal a significant relationship between the built environment and physiological as well as psychological states. For instance, thermal comfort was found to be the most commonly studied IEQ variable, affecting heart activity and skin temperature. The research also identifies the need for a shift towards using advanced technologies like Mobile Brain/Body Imaging (MoBI) for capturing real-time physiological data in natural settings.

Major conclusions include the need for a multi-level, evidence-based approach that considers the dynamic interaction between the brain, body, and environment. The study advocates for the incorporation of multiple physiological signals to gain a comprehensive understanding of well-being in relation to the built environment. It also highlights gaps in current research, such as the absence of noise as a studied variable of IEQ and the need for standardized well-being assessment tools. By synthesizing these insights, the research aims to pave the way for future studies that can inform better design and policy decisions for indoor environments.

## Introduction

The built environment can be defined as a natural environment that has been modified by human conceptualizations and actions (Bartuska, 2007; Portella, 2014). Even when the built environment is not a synonym for the urban environment (i.e., rural areas also incorporate modifications made by human actions (Alanazy et al., 2019); *urban* has been defined as a transformed natural environment that incorporates nonagricultural elements of population density and social-economic organization (Weeks, 2010). Given the characteristics of urban areas, such as job opportunities, and access to health, social, and educational services (Bodo, 2019; Hall et al., 2006; McMichael, 2000; Pateman, 2011; Weeks, 2010), more than 50% of the world’s population lives in urban places (Ritchie & Roser, 2018; The World Bank, 2022; United Nations, 2018), a figure expected to rise to 68% by 2050 (Ritchie & Roser, 2018; The World Bank, 2022).

People not only live in more urbanized spaces but also spend the majority of their lives (about 90%) *inside* the built environment; this means inside human-made constructions (Azzazy et al., 2021; Klepeis et al., 2001; Schweizer et al., 2007). Considering this peculiarity of modern life, researchers have explored several components of the built environment (e.g., indoor characteristics, access to green spaces and blue views, light exposure, and air characteristics) to understand its relationship with physiology (e.g., cortisol level, circadian rhythm; (Beil & Hanes, 2013; Stevens & Rea, 2001), psychological states (e.g., happiness, irritability, stress; Sullivan & Chang, 2011; Zainal & Hosni, 2022), and cognitive processes (e.g., attention, learning, memory; Besser et al., 2018; Keis et al., 2014; Marchand et al., 2014; Möystad, 2017). Findings show that some features of the environment can promote or disturb mental health and cognitive functioning (Karakas & Yildiz, 2020; Moore et al., 2018; Rhodes et al., 2018), and that can have a direct impact on neurobiology (Djebbara et al., 2022).

### Built environment’s impact on health

Several studies have shown that crowded, noisy, and perceived-as-dangerous places might negatively impact psychological states, for instance, by causing chronic stress states that are associated with psychiatric symptoms (Evans et al., 2003; Lederbogen et al., 2011, 2013; Matthews & Yang, 2010). For example, traffic-related pollution has been correlated with negative mental health outcomes such as increased anxiety and depressive symptoms (Beemer et al., 2021; Pelgrims et al., 2021; Power et al., 2015).

In contrast, other characteristics have been shown to positively impact psychological states. Exposure to green areas might significantly alleviate mental fatigue and restore calm from stressful conditions, contributing to healthy psychological development in children (Sullivan & Chang, 2011). Places with green views or direct access to trees or other vegetation have been associated with increased perceived well-being (Day, 2008; Sullivan & Chang, 2011; Yuen & Nyuk Hien, 2005).

Similarly, natural light (i.e., daylight) exposure might reduce the intensity of symptoms of seasonal depression (Elliott et al., 1993; Matthews & Yang, 2010; Taylor et al., 1997). Overall, the increase in environmental quality has proven to reduce both physiological and psychological negative responses in workplaces, hospitals, and even in artificial simulations (Beil & Hanes, 2013; Beukeboom et al., 2012; Codinhoto et al., 2009; Lottrup et al., 2013; Ulrich, 1981; van den Berg et al., 2010; Ward Thompson et al., 2012).

Regarding indoor environments, several characteristics have been explored due to their relation to mental health and well-being (Rohde et al., 2020). The study of these characteristics has been conceptualized under the idea of Indoor Environmental Quality (IEQ). IEQ is a set of conditions for indoor environments and their impact on inhabitants’ health and well-being (Abdulaali et al., 2020; Mallawaarachchi et al., 2012; Steinemann et al., 2017). The results of some systematic literature reviews show that some of the most common positive IEQ conditions are air quality, ergonomics, and thermal, acoustic, and visual comfort (Al horr et al., 2016; Salonen et al., 2013).

Hence, these conditions have been explored in different environments and populations. For example, Turunen and colleagues (2014) studied the characteristics of classroom environments and their impact on student’s health and well-being. They found a relation between symptom presentation in 6th-grade students, such as fatigue, headache, and stuffy nose, with environmental variables like acoustic contamination (i.e. noise) and poor ventilation. Also, the review conducted by Salonen and colleagues (2013) shows that indoor environmental factors (e.g., acoustic environment, ventilation, visual conditions) can have a positive effect or hinder the health and well-being of users of healthcare facilities (e.g., patients, medical staff, relatives).

### A new perspective for neuroscience research of the built environment

An alternative paradigm to classic conceptualizations in cognitive science conceives the nature of the cognitive process as a complex phenomenon that emerges from the dynamic relationship between the brain/body system of an agent in active interaction with its environment (Jonas, 1966; Newen et al., 2018; Thelen & Smith, 1995; Varela et al., 1991). Under this paradigm, cognition is understood as an embodied, environmentally scaffolded, and enactive process. This means: 1) that the brain and all the agent’s biology play a major role in cognition (Thompson, 2010; Varela et al., 1991); that cognitive processes are dependent on the environment and also have been transformed by environmental resources (Laland et al., 2000; Stephan & Walter, 2020; Sterelny, 2010); and 3) the mind is the product of the dynamic relationship between brain/body and environment (Clark, 2000; Thompson, 2010). Hence, the consideration of intracranial (e.g., brain activity) and extracranial dynamics (e.g., body, environment) allows the understanding of cognitive processes as embodied in biology and scaffolded by the environment (De Jaegher et al., 2010; Di Paolo et al., 2010; Di Paolo & De Jaegher, 2012; Kyselo, 2014; Laland et al., 2000; Parada & Rossi, 2018; Rossi et al., 2019; Sterelny, 2010). This paradigm to understand and study cognition has been conceptualized as the 3E cognition perspective or 3E-Cognition^1^. These new ideas allow the development of new research questions and hypotheses about the relationships between the body, brain, mind, and environment (S. Lee et al., 2022; Varela et al., 1991). Hence, this perspective can be considered an effort to generate a multilevel evidence-based approach for studying this dynamic system composed of the brain and the body in its interaction with the environment and its implication for cognition, psychological status, and even social interactions.

Considering the aforementioned concepts, questions about the impact of different components of the built environment on physiology are raised. This field of research has primarily addressed body temperature and brain and heart activity. For instance, the impact of thermal characteristics of the environment on the body has been explored in previous research due to the connection between vagal and sympathetic nerve activity and its role in thermoregulation (Yao et al., 2009). Hence, thermal comfort has been studied in relation to its impact on heart activity, showing that a higher ratio of low-frequency/high-frequency relates to unpleasant thermal sensation and discomfort (Liu et al., 2008; Persiani et al., 2021; Zhu et al., 2018). Similarly, Mulders and colleagues (2020) assessed brain activity concerning thermal stimuli, showing differences in terms of time-frequency analyses.

Furthermore, alpha activity during exposure to natural settings presented similar characteristics to alpha activity during relaxation and restoration states regarding alpha-theta oscillations and synchronization (Bolouki, 2022; Chen et al., 2020).

### Methodological implications for neuroscience research

Research from the neuroscience field has incipiently addressed the relationship between environmental variables, cognitive processes, and physiological states using different methods. For instance, some studies have utilized real-time biometric data (e.g., electrodermal activity, EDA; heart rate, HR) as a measure of physiological reactions to the built environment (Kim et al., 2019; Persiani et al., 2021); electric brain activity (e.g., via electroencephalogram, EEG) to assess the impact of the built environment attributes on brain activity (Banaei et al., 2017; Cheng et al., 2022; M. Hu & Roberts, 2020; Jiang et al., 2020), and the implementation of self-report measured as the most common technique to collect information from participants (Azzazy et al., 2021; Jiang et al., 2020; Yadav et al., 2018).

As shown by Azzazy and colleagues (2021), the study of brain activity has remained to classical paradigms (e.g., picture presentations; (Jiang et al., 2020), and non-mobile techniques for data acquisition (e.g., functional magnetic resonance imaging, fMRI; computerized tomography, CT). Only during the past decade researchers have changed paradigms to study brain dynamics in the built environment using virtual reality (Cheng et al., 2022; M. Hu & Roberts, 2020; Li et al., 2020) or in real-world settings (Azzazy et al., 2021; Banaei et al., 2017, 2020; Mavros et al., 2016, 2022; Neale et al., 2020).

These findings are consistent with new technological advancements of the last 15 years related to the Mobile Brain/Body Imaging (MoBI) framework, which allows measuring different body signals during natural movement (Gramann et al., 2011, 2014; Jungnickel & Gramann, 2016; Makeig et al., 2009). This technical-methodological approach has increased the possibility of studying physiological variables in ecologically valid paradigms and real-world situations as they naturally unfold (Gramann et al., 2014; Ladouce et al., 2016; S. Lee et al., 2022; Matusz et al., 2019; Parada & Rossi, 2021; Rojas-Líbano & Parada, 2019). By doing so, it is possible to embrace the complexity of the relationship between the body, mind, and environment and address a broader understanding of their relationship and physiological dynamics as they naturally unfold in interaction with the built environment.

### The present review

This review will focus on the IEQ variables since they have been defined as a proxy for environmental quality and comfort, which are closely related to well-being. Previous reviews have already addressed the relationship between the built environment and well-being from a neuroscientific perspective (Ancora et al., 2022; Azzazy et al., 2021; S. Lee et al., 2022). However, the present review is situated from a 3E perspective, so it aims to incorporate not only brain activity but multiple physiological signals in the study of the impact of the indoor built environment on psychological well-being, as well as a technical-methodological perspective for analyzing the existent literature.

In this sense, the present review aims to systematize the current scientific evidence on methodologies used to investigate the impact of indoor built environment on well-being, focusing on assessing physiological variables. By doing so, we intend to 1) categorize the main IEQ variables studied in research investigating the impact of the built indoor environment on well-being in real-life settings; 2) identify and summarize methodological aspects (e.g., data collection techniques, study settings, and biomarkers) used to explore the relationship between the indoor built environment and well-being; and 3) review the self-report instruments employed to capture subjective experiences related to indoor environmental quality and well-being.

## Methodology

### Protocol and Registration

This systematic review was conducted following the Joanna Briggs Institute (JBI) guidelines for systematic reviews (Aromataris & (Eds.)., 2020; Santos et al., 2018) and following the Preferred Reporting Items for Systematic Reviews and Meta-Analyses (PRISMA) guidelines (Page et al., 2021). The protocol can be found at Open Science Framework (OSF)^2^.

### Eligibility Criteria

Empirical studies on indoor built environment quality incorporating physiological and well-being self-report measurements were assessed for eligibility. To be included in the present review, studies should: 1) Have neurotypical human participants; 2) Study one or more of the four variables related to Indoor Environmental Quality (i.e., air quality, thermal comfort, noise, lightning); Be an empirical article that addresses the relation with, at least, one physiological signal; studies only incorporating behavioral measures (e.g., only implementing eye tracking or motion energy analysis) were not considered for eligibility since the main purpose of this review is to assess physiological variables; 4) Incorporate at least one self-reported measure of well-being; and 5) Be conducted in real-world scenarios instead of laboratory setups (e.g., work or residential environments).

All experimental contexts were included in this review. All available publications in English or Spanish were eligible for inclusion, with no limitation regarding time. All observational and experimental designs were considered, and no gray literature (i.e., reports that haven’t been included in a peer review process) was included. Table 1 provides an overview of the inclusion and exclusion criteria applied to the articles.

**Table 1.** PICO framework for inclusion and exclusion criteria.

| **PICO** | **Inclusion criteria** | **Exclusion criteria** |
| --- | --- | --- |
| Population | Neurotypical human participants | None |
| Intervention | Exposure to at least one of the four main factors of Indoor Environmental Quality—air quality, thermal comfort, noise, or lighting—in real-world scenarios instead of traditional laboratory setups | None |
| Comparators | Not applicable | Not applicable |
| Outcomes | Data related to acquisition of physiological variables and self-reported measures of well-being | None |
| Other | Empirical design, published in peer-reviewed journals, available in English or Spanish, from any geographic region, with no time restrictions. | None |

### Information Sources

For this review, we conducted a comprehensive literature search on electronic databases in Web of Science (WOS), PUBMED, and SCOPUS. The reference list of key articles and reviews was screened for additional studies (Azzazy et al., 2021; Beemer et al., 2021; S. Lee et al., 2022; Yeo, 2021). All databases were consulted during October 2024.

### Search

A search strategy was developed following the Peer Review of Electronic Search Strategies (PRESS) checklist (McGowan et al., 2016). The search strategy was adapted to each database. The search process was conducted by two members of the research team^3^ (AGC and MAP) after a preliminary, non-systematic review of keywords related to indoor environmental quality. The keywords for searching articles were related to indoor environmental quality (e.g., thermal comfort, noise) and referred to specific data acquisition techniques (e.g., electroencephalogram) and/or physiological signal type (e.g., brain activity). Keywords were searched in the articles’ abstracts.

Systematics reviews and conference abstracts were excluded from the search.

### Selection of Sources of Evidence

The search process results were exported into Microsoft Excel (Microsoft Corporation, 2018), and all duplicated articles were removed. Four independent reviewers (AGC, MAP, EV, and JMS) performed the screening process for article inclusion. First, the reviewers selected articles based on the articles’ titles and then based on abstract information. For each stage, a consistency analysis was conducted to determine the level of agreement regarding eligibility criteria. After achieving ∼90% of agreement, each reviewer selected articles independently. Disagreements between reviewers were addressed through multiple rounds of discussion.

### Data Charting Process

The reviewer team developed a Google form questionnaire for the data charting process. The questionnaire asked questions about article descriptions and items inspired by the present review’s goals. To ensure internal consistency, four authors (AGC, MAP, EV, and JMS) codified the first 10 articles, and the rest were divided equally to be reviewed independently by the same researchers.

### Data Items

The questionnaire’s items were referred to: 1) the article’s characterization (e.g., year of publication, country), and 2) the methodological considerations (e.g., aim, environmental variable, data collection technique, setting, and task, among others).

### Risk of Bias

The potential risk of bias in the studies was assessed using the JBI Critical Appraisal Tools (Moola et al., 2017; Munn et al., 2020), with the specific tool extension applied based on the study design of each included article. Two authors conducted this process independently (MAP and EV).

### Synthesis of Results

Following a narrative methodology that Arksey and O’Malley (2005) presented, data were summarized following the data extraction categories to answer the review objectives and questions. Data will be presented graphically (graphs or diagrams) and in tables.

## Results

### Study selection

After conducting the search process in all three databases, 2.701 articles were found. After removing duplicated articles, 1.562 articles were left. To select the articles, we conducted a two-stage screening process. Based on the information presented in the title and abstract, 1.189 articles were excluded. After the full-text review, a total of 16 articles were included in the present review.

Among the principal reasons for exclusion are: 1) Only use of self-report measures to study an IEQ variable (N = 117); 2) Studying a non-target characteristic of Indoor Environmental Quality (N = 79); 3) Only use of behavioral measures (N = 59); and 4) Study conducted in traditional laboratory settings (N = 49).

Other 12 articles were found manually using existing reviews’ citations. These reports were excluded: eight articles were duplicates, and four focused on non-target characteristics of indoor environmental quality. Figure 1 shows the flow of articles across the different stages.

**Figure 1.**
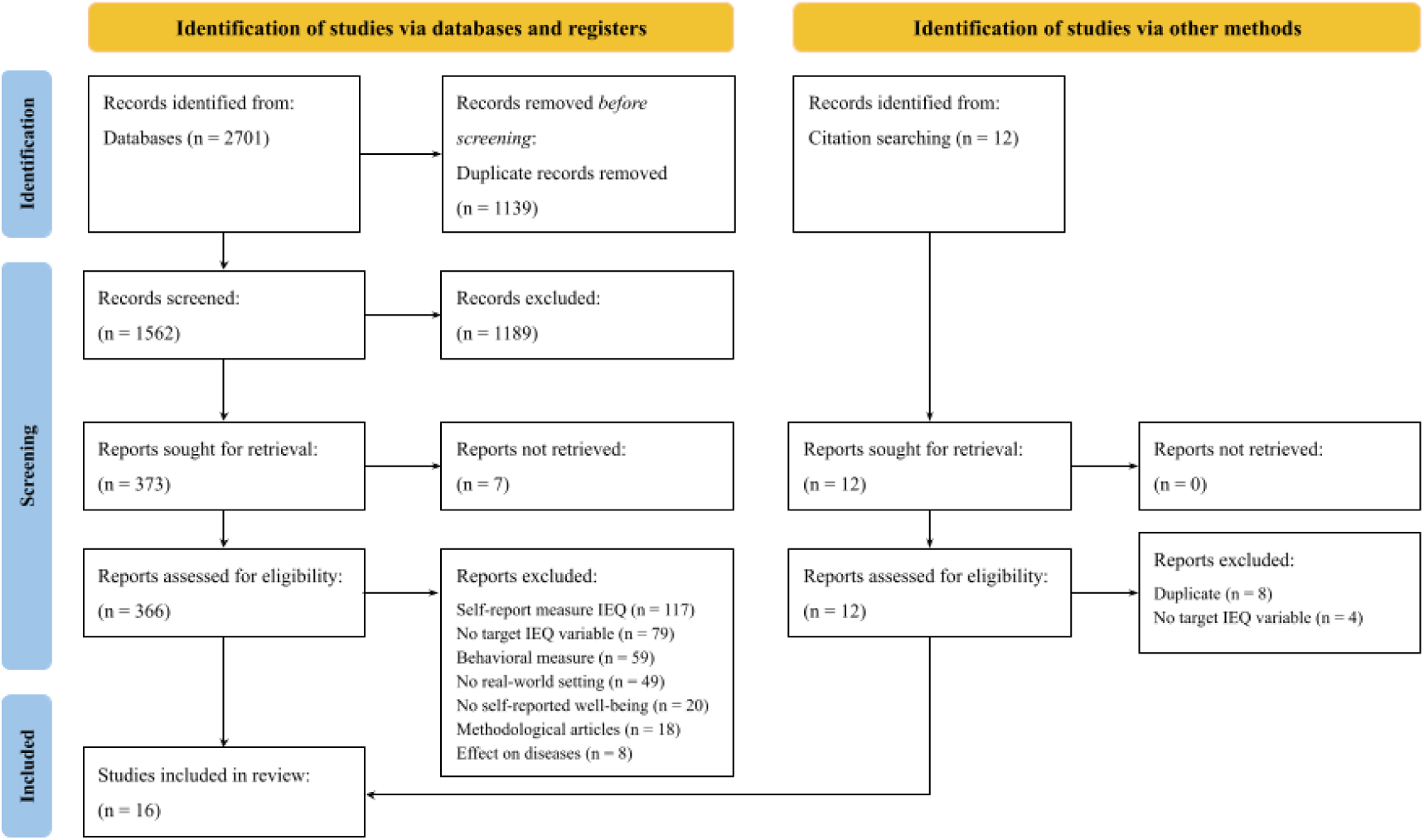
PRISMA flow diagram

### Study characteristics

The publication dates of the included articles ranged from 2017 to 2024. The results show that 11 out of 18 articles (69%) were conducted in the last four years (Figure 2). Regarding geographical location, there is a predominant presence of studies conducted in the Asian continent, with 8 of them performed in China (Fanpu et al., 2024; Gao et al., 2023; J. Hu et al., 2022; Song et al., 2020; Wang et al., 2023; Wu & Wagner, 2023; Zhou et al., 2023; Zhu et al., 2018). Switzerland follows with 2 articles (Chinazzo et al., 2018, 2019), while the remaining articles were published in various other countries, including Italy (Barbic et al., 2022), USA (Beaudette et al., 2024), Sweden (Fischl & Johansson, 2024), Japan (Okamoto et al., 2017), India (Roy et al., 2024), and England (Snow et al., 2019). Interestingly, all included studies were conducted in the Northern Hemisphere, showing a lack of research from southern parts of the world (Figure 3).

**Figure 2.**
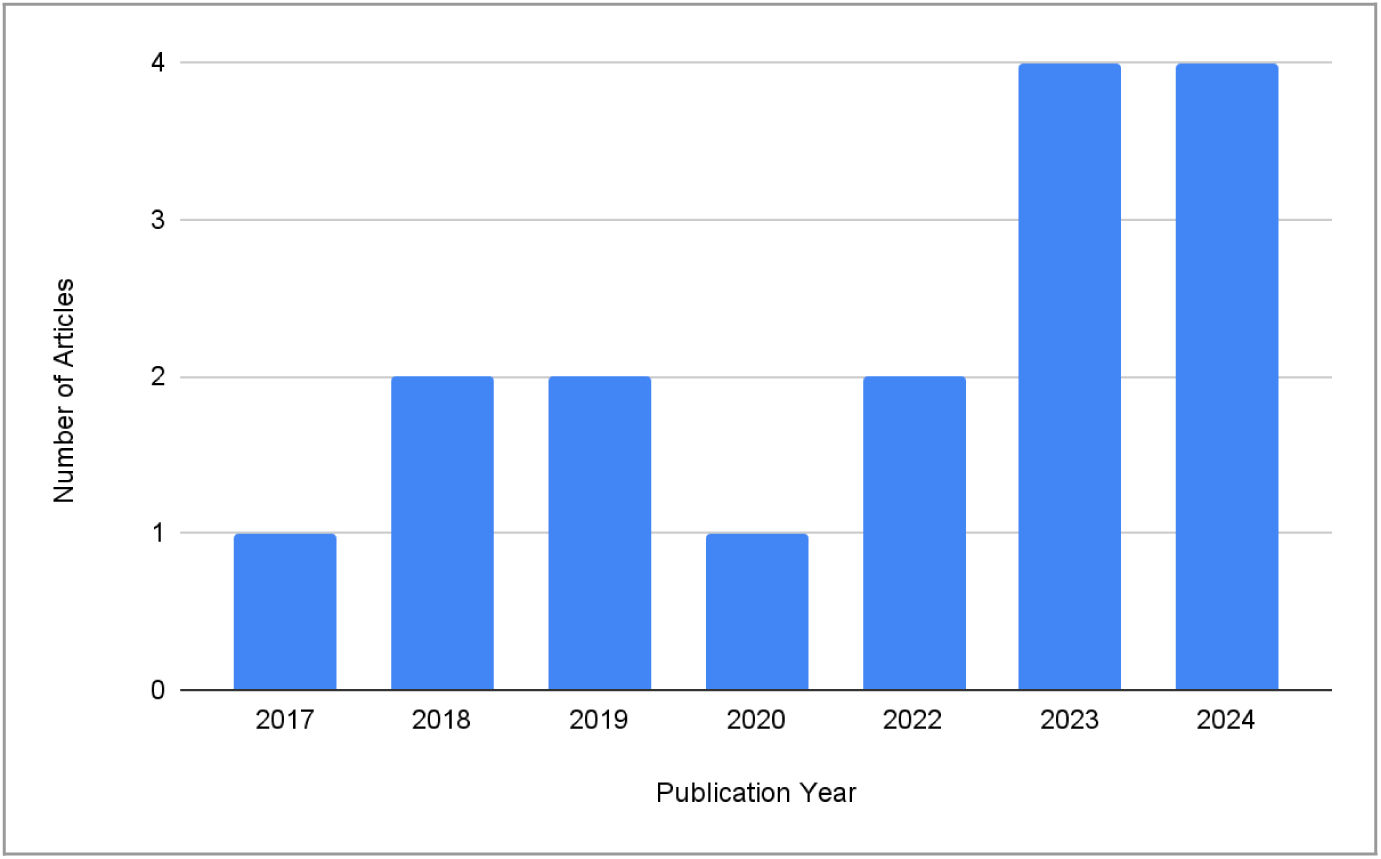
Distribution of included articles by year of publication

**Figure 3.**
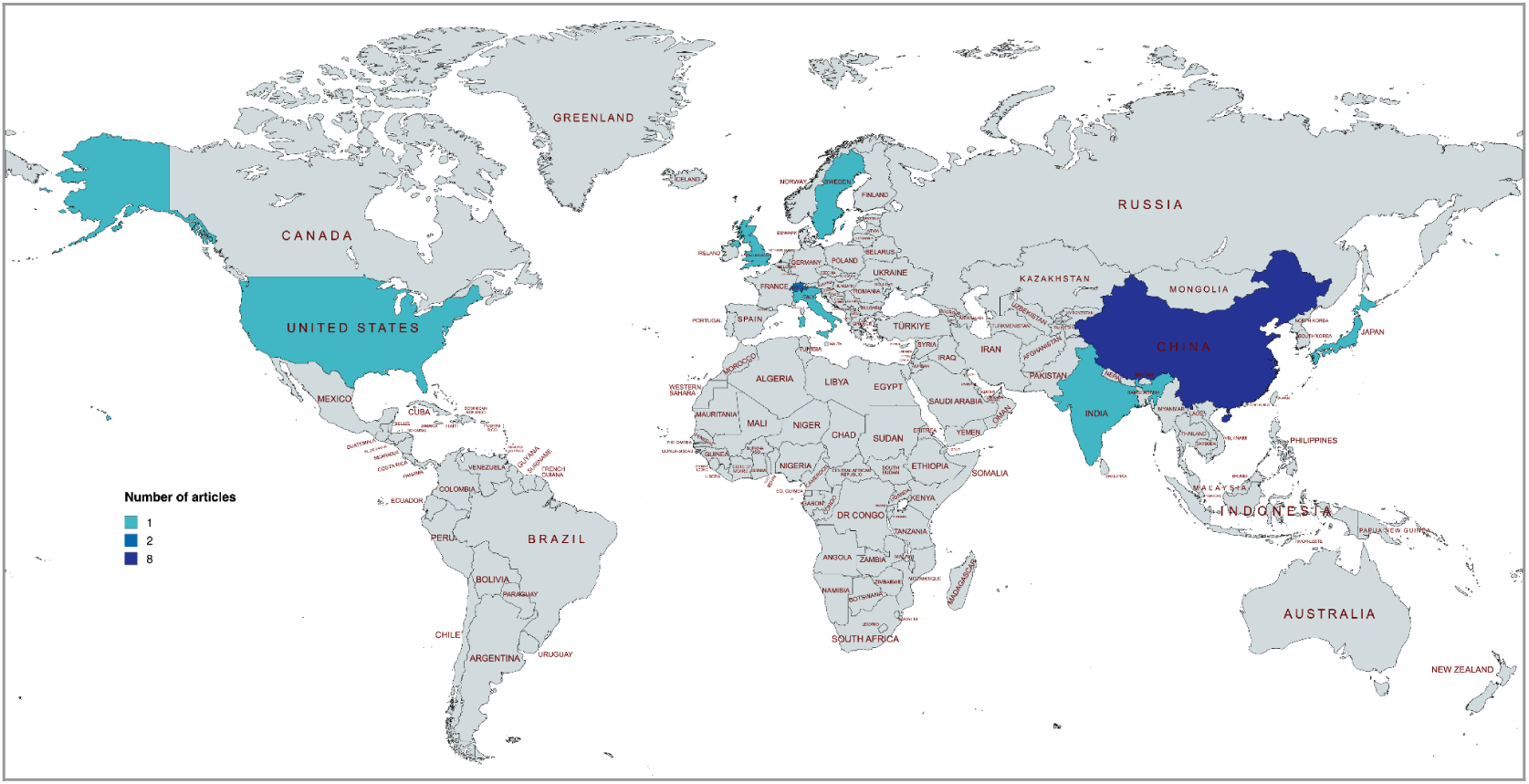
Geographical distribution of the included articles

### Risk of Bias

The results of the risk of bias assessment are summarized in Table 2. Each study included in this review was evaluated using the design-specific JBI Critical Appraisal Tool. Of the 14 case series studies, 13 met most of the criteria outlined by the tool. However, it was unclear whether participants were consecutively included in all of these studies.

**Table 2.** Summary of risk of bias.

| **Case Series Studies<sup>4</sup>** |  |  |  |  |  |  |  |  |  |  |
| --- | --- | --- | --- | --- | --- | --- | --- | --- | --- | --- |
| **Reference** | **Q1** | **Q2** | **Q3** | **Q4** | **Q5** | **Q6** | **Q7** | **Q8** | **Q9** | **Q10** |
| Okamoto et al., 2017 | Yes | Yes | Yes | Unclear | Unclear | Yes | NA | Yes | No | Yes |
| Wang et al., 2018 | Yes | Yes | Yes | Unclear | Yes | Yes | NA | Yes | Yes | Yes |
| Chinazzo et al., 2018 | Yes | Yes | Yes | Unclear | Yes | Yes | NA | Yes | Yes | Yes |
| Snow et al., 2019 | Yes | Yes | Yes | Unclear | Yes | Yes | NA | Yes | Yes | Yes |
| Chinazzo et al., 2019 | Yes | Yes | Yes | Unclear | Yes | Yes | NA | Yes | Yes | Yes |
| Song et al., 2020 | Yes | Yes | Yes | Unclear | Yes | Yes | NA | Yes | Yes | Yes |
| Barbic et al., 2022 | Yes | Yes | Yes | Unclear | Yes | Yes | NA | Yes | Yes | Yes |
| Wang et al., 2023 | Yes | Yes | Yes | Unclear | Yes | Yes | NA | Yes | Yes | Yes |
| Zhou et al., 2023 | Yes | Yes | Yes | Unclear | Yes | Yes | NA | Yes | Yes | Yes |
| Gao et al., 2023 | Yes | Yes | Yes | Unclear | Yes | Yes | NA | Yes | Yes | Yes |
| Roy et al., 2024 | Yes | Yes | Yes | Unclear | Yes | Yes | NA | Yes | Yes | Yes |
| Fischl & Johansson, 2024 | Yes | Yes | Yes | Unclear | Yes | Yes | NA | Yes | Yes | Yes |
| Fanpu et al., 2024 | Yes | Yes | Yes | Unclear | Yes | Yes | NA | Yes | Yes | Yes |
| Beaudette et al., 2024 | Yes | Yes | Yes | Unclear | Yes | Yes | NA | Yes | Yes | Yes |
| **Cross-Sectional Studies<sup>5</sup>** |  |  |  |  |  |  |  |  |  |  |
| **Reference** | **Q1** | **Q2** | **Q3** | **Q4** | **Q5** | **Q6** | **Q7** | **Q8** |  |  |
| Hu et al., 2022 | Yes | Yes | Yes | Yes | Unclear | Unclear | Yes | Yes |  |  |
| Wu & Wagner, 2023 | Yes | Yes | Yes | Yes | Unclear | Unclear | Yes | Yes |  |  |

*NA: not applicable*

Regarding the cross-sectional studies, both met 6 out of the 8 criteria analyzed. However, the articles lacked sufficient information to clearly address items related to identifying and managing potential confounders.

### Synthesis of Results

#### Aim

Regarding the main aim of the articles included in this review, it is possible to notice a diversity of objectives pursued. This mainly refers to the development of different hypotheses regarding the impact of the indoor built environment on physiology and well-being/cognition. Despite this diversity, the main objective is to study the effects of aspects of the indoor built environment on physiology and cognition. Table 3 displays the main objective of each included article in this review.

**Table 3.** Main objectives and IEQ variables studied in the included articles.

| Reference | Aim | IEQ Variable |
| --- | --- | --- |
| Okamoto et al., 2017 | To determine the effect of airflow on brain activity using electroencephalograms (EEG) of participants in a living environment under different airflow conditions. | Thermal comfort |
| Chinazzo et al., 2018 | To investigate the effect of daylight transmitted through three colored glazing types on thermal responses and overall comfort at three temperature levels. | Thermal comfort |
| Wang et al., 2018 | To study the effect of the indoor environment on students' learning performance in summer. | Thermal comfort |
| Chinazzo et al., 2019 | To investigate the influence of daylight on thermal responses, intended as both subjective thermal perceptions and physiological responses, by studying different combinations of daylight and temperature levels. | Lighting, Thermal comfort |
| Snow et al., 2019 | To measure changes in human performance, physiological, neurophysiological (EEG), and psychological factors as a result of elevated indoor CO<sub>2</sub> concentrations; and to validate EEG as an objective measurement of sleepiness for studies concerned with the effect of the indoor environment on humans. | Air quality |
| Song et al., 2020 | To identify the important local thermal conditions for sleeping thermal comfort. | Thermal comfort |
| Barbic et al., 2022 | To evaluate the effects of reducing classroom temperatures on cognitive performance and to evaluate the associated changes in cardiac autonomic control in a class of undergraduate students. | Thermal comfort, Air quality |
| Hu et al., 2022 | To analyze gender differences in thermal comfort, work performance, and Sick Building Syndrome symptoms and to define optimal temperature ranges considering gender differences in the three above-mentioned aspects in classrooms in winter. | Thermal comfort |
| Gao et al., 2023 | To analyze the dynamic changes and effects of indoor temperature and exercise behavior on human thermal comfort and establish an evaluation mechanism of exercise thermal comfort. | Thermal comfort |
| Wang et al., 2023 | To provide theoretical foundations for developing healthy building Smell-scapes with essential oil. | Thermal comfort, Lighting, Air quality |
| Wu & Wagner, 2023 | To examine the effect of outdoor short-term thermal history on the thermal comfort and physiological responses of humans in the indoor environment. | Thermal comfort |
| Zhou et al., 2023 | To determine the preferred indoor air velocity under hot and humid conditions and to examine the effects of elevated air movement on subjective perceptions and skin temperature. | Thermal comfort |
| Beaudette et al., 2024 | To investigate the use of readily available heating devices in typical indoor climate-controlled environments: contexts in which some users may feel uncomfortably cool or cold. | Thermal comfort |
| Fanpu et al., 2024 | To explore the best artificial illumination supplement under the comprehensive evaluation of vision, emotion, and cognition in different periods. | Lighting |
| Fischl & Johansson, 2024 | To integrate digital occupancy assessment methods to understand indoor lighting conditions' impact on occupant well-being. | Lighting |
| Roy et al., 2024 | To explore the influence of three different lighting conditions on participants' perceptions of task lighting attributes, aspects of room aesthetics, and their physiological responses. | Lighting |

#### Indoor Environmental Quality Variable

The IEQ construct involves four main dimensions: Thermal Comfort, Air Quality, Noise, and Lightning. As shown in Table 3, this review identified Thermal Comfort as the most common dimension studied (N = 12; Barbic et al., 2022; Beaudette et al., 2024; Chinazzo et al., 2018, 2019; Gao et al., 2023; Hu et al., 2022; Okamoto et al., 2017; Song et al., 2020; Wang et al., 2018; Wang et al., 2023; Wu & Wagner, 2023; Zhou et al., 2023), followed by Lighting (N = 5; Chinazzo et al., 2019; Fanpu et al., 2024, 2024; Fischl & Johansson, 2024; Roy et al., 2024; Wang et al., 2023) and Air Quality (N = 3; Barbic et al., 2022; Snow et al., 2019; Wang et al., 2023). Noise was not explicitly addressed in the studies reviewed, highlighting a gap in research on this dimension.

Only 3 studies (18,75%) considered more than one variable of the IEQ constructs in their research questions (Barbic et al., 2022; Chinazzo et al., 2019; Wang et al., 2023), emphasizing a tendency to focus on isolated aspects of the indoor environment rather than adopting a holistic perspective. Notably, these three studies consider thermal comfort along with another IEQ factor.

#### Task

The reviewed articles described a diversity of tasks during environmental manipulation and physiological recording. The most common types of tasks consisted of office-related tasks or similar activities, such as reading or drawing (N = 8; Beaudette et al., 2024; Chinazzo et al., 2018, 2019; Fanpu et al., 2024; Fischl & Johansson, 2024; Hu et al., 2022; Roy et al., 2024; Zhou et al., 2023). Neuropsychological assessments, like the Stroop test, were also frequently used (N = 5; Barbic et al., 2022; Okamoto et al., 2017; Snow et al., 2019; Wang et al., 2018; Wang et al., 2023). Less common tasks included physical activity (N = 1; Gao et al., 2023) and sleeping (N = 1; Song et al., 2020).

Additionally, one study did not specify the nature of the task performed by participants (Wu & Wagner, 2023). Table 4 summarizes the tasks performed during measurements in the included articles.

**Table 4.** Task performed and research settings.

| Reference | Task | Setting |
| --- | --- | --- |
| Okamoto et al., 2017 | Time-perception task | Show house |
| Chinazzo et al., 2018 | Office tasks (paper-based performance tests such as the d2 test) | Office-like room |
| Wang et al., 2018 | Learning performance test | Experimental room in rural primary and secondary school |
| Chinazzo et al., 2019 | Office tasks (assigned paper-based) | Office-like room |
| Snow et al., 2019 | Cognitive test (Stroop Test, Shifting Attention Task, Continuous Performance Test, Four Part Continuous Performance Test) | Office |
| Song et al., 2020 | Sleeping behavior | Residential house |
| Barbic et al., 2022 | Cambridge Brain Science Task | University classroom |
| Hu et al., 2022 | Course-related activities such as reading books or doing homework | University classroom |
| Gao et al., 2023 | Field test lasting >100 min (preparation, warm-up and testing stage). | Physical fitness training center |
| Wang et al., 2023 | Cognitive Tasks/Tests (Stroop, N-Back, Visual Search Task, Digital Memory Span Test) | Office |
| Wu & Wagner, 2023 | No specific task (indoor staying) | University dormitory rooms |
| Zhou et al., 2023 | Office tasks (computer work or reading) | Office-like rooms |
| Beaudette et al., 2024 | Office tasks (self-directed desk work) | Office-like room |
| Fanpu et al., 2024 | Reading tasks (Anfimov letter recognition table, number proofreading table, and Randall ring checklist) | Reading space of the university library |
| Fischl & Johansson, 2024 | Simple (navigation map) and complex (floor plans and facades of their homes) drawing tasks | Students group rooms |
| Roy et al., 2024 | Reading (common textbook) and writing (copy diagrams and paragraphs) tasks | Tertiary educational institute classroom |

#### Setting/Context

This review focused exclusively on studies conducted in real-life contexts (Table 4). The most common settings were educational or university environments (N = 7; Barbic et al., 2022; Fanpu et al., 2024; Fischl & Johansson, 2024; Hu et al., 2022; Roy et al., 2024; Wang et al., 2018; Wu & Wagner, 2023), followed by professional or workplace settings, specifically offices and office-like environments (N = 6; Beaudette et al., 2024; Chinazzo et al., 2018, 2019; Snow et al., 2019; Wang et al., 2023; Zhou et al., 2023). Additionally, residential contexts (N = 2; Okamoto et al., 2017; Song et al., 2020) and a physical training center (N = 1; Gao et al., 2023) were included.

#### Data Collection Technique

The reviewed articles employed various data collection techniques, depending on the physiological variables of interest. For cardiac activity, the most common methods included heart rate monitors (N = 6; Barbic et al., 2022; Chinazzo et al., 2018; Fanpu et al., 2024; Fischl & Johansson, 2024; Gao et al., 2023; Wang et al., 2023), the use of sphygmomanometers (N = 3; Roy et al., 2024; Wang et al., 2018; Wang et al., 2023), pulse oximeters (N = 3; Hu et al., 2022; Roy et al., 2024; Snow et al., 2019) and blood pressure monitors (N = 2; Gao et al., 2023; Wu & Wagner, 2023). Regarding skin temperature, surface thermometers were the most frequently used devices (N = 6; Beaudette et al., 2024; Gao et al., 2023; Hu et al., 2022; Snow et al., 2019; Wang et al., 2018; Wu & Wagner, 2023), followed by iButtons (N = 4; Chinazzo et al., 2018, 2019; Song et al., 2020; Zhou et al., 2023) and thermographers (N = 2; Gao et al., 2023; Okamoto et al., 2017). Only 2 articles collected information on electrical brain activity by using electroencephalography (EEG; Okamoto et al., 2017; Snow et al., 2019). Table 5 summarizes the instruments used and measurements conducted in the included articles.

**Table 5.** Instruments used and measurements conducted.

| Reference | Data collection technique | Main biomarker | Self-report |
| --- | --- | --- | --- |
| Okamoto et al., 2017 | EEG; Thermographer | EEG frequency bands; Skin temperature | Predicted Mean Vote (PMV); Predicted Percentage of Dissatisfied (PPD) |
| Chinazzo et al., 2018 | iButton; Wristband sensor | Skin temperature; Heart rate; Skin conductance | Thermal comfort (based on 7-point scale of the ASHRAE); Ergonomics of the thermal environment |
| Wang et al., 2018 | Electronic sphygmomanometer; Electronic thermometer | Blood pressure; Heart rate; Body temperature | ASHRAE 7-points thermal sensation scale |
| Chinazzo et al., 2019 | iButton | Skin temperature | Ergonomics of the thermal environment |
| Snow et al., 2019 | EEG; Electrodes; Finger clip; Abdominal belt | Heart rate; Respiration rate; Skin temperature; EEG frequency bands | Sick Building Syndrome symptoms; Stanford Sleepiness Scale; Positive and Negative Affective State |
| Song et al., 2020 | iButton | Skin temperature | Thermal sensation, comfort, and acceptability evaluation scales (TSV; TCV; TAV) |
| Barbic et al., 2022 | ECG | Heart rate; Respiration rate | Questionnaire for the Thermal Comfort Survey (modified by Wang) |
| Hu et al., 2022 | Thermocouple; Fingertip pulse oximeter | Skin temperature; Blood oxygen saturation; Heart rate | Thermal comfort; Self-estimated work performance; Sick Building Syndrome symptoms (based on 7-point scale of the ASHRAE) |
| Gao et al., 2023 | Infrared imager; Four channels thermometer; Chest strap sensor; Blood pressure monitor | Skin temperature; Heart rate; Blood pressure | TSV; TCV; TPV; TSL; FSV |
| Wang et al., 2023 | Holter Monitor; Sphygmomanometer | Heart rate; Heart rate variability; Blood pressure | POMS questionnaire; Indoor air quality acceptability scale; Odor scale |
| Wu & Wagner, 2023 | Surface thermometer; Blood pressure monitor | Skin temperature; Heart rate; Blood pressure | Thermal sensation, preference, comfort, and acceptability (based on 7-point scale of the ASHRAE) |
| Zhou et al., 2023 | iButton | Skin temperature | Scales for perceptions on temperature, humidity, and air movement (TSV; HSV; AMV; TCV; TS; HS; AS; TAV; HAV; AMA; TPV; HPV; AVP) |
| Beaudette et al., 2024 | Thermistors | Skin temperature | Adapted ASHRAE scale (perceived thermal sensation and comfort) |
| Fanpu et al., 2024 | Heart rate band | Heart rate variability | Positive and Negative Emotion Scale; Visual discomfort scale; Office lighting survey scale; Subjective alertness scale |
| Fischl & Johansson, 2024 | Galvanic skin response sensor; Wristband sensor | Skin conductance, Heart rate | 7-point Likert scale (satisfaction, pleasantness, novelty, and other perceptions); Self-Assessment Manikin (emotional reaction) |
| Roy et al., 2024 | Digital sphygmomanometer, Fingertip pulse oximeter | Blood pressure, Heart rate | 5-point semantic differential scale (aspects of lighting of tasks and room aesthetics) |

*TSV: thermal sensation vote; TCV: thermal comfort vote; TAV: thermal acceptability vote; TPV: thermal preference vote; TSL, thermal satisfaction level; FSV, fatigue sensation vote; POMS: profile of mood states; HSV: humidity sensation vote; AMV: air movement sensation vote; TS: thermal satisfaction; HS: humidity satisfaction; AS: air movement satisfaction; TAV: thermal acceptability vote; HAV: humidity acceptability vote; AMA: air movement acceptability vote; HPV: humidity preference vote; AVP: air movement preference vote.*

#### Biomarker

In the reviewed articles, the analyzed biomarker was restricted to the type of physiological signal collected (Table 5). The most common units of analysis for cardiac activity were heart rate (N = 10; Barbic et al., 2022; Chinazzo et al., 2018; Fischl & Johansson, 2024; Gao et al., 2023; Hu et al., 2022; Roy et al., 2024; Snow et al., 2019; Wang et al., 2018; Wang et al., 2023; Wu & Wagner, 2023) and blood pressure (N = 5; Gao et al., 2023; Roy et al., 2024; Wang et al., 2018; Wang et al., 2023; Wu & Wagner, 2023). For temperature-related measurements, the main biomarker was skin temperature (N = 10; Beaudette et al., 2024; Chinazzo et al., 2018, 2019; Gao et al., 2023; Hu et al., 2022; Okamoto et al., 2017; Snow et al., 2019; Song et al., 2020; Wu & Wagner, 2023; Zhou et al., 2023). The most frequently analyzed parameter regarding respiratory activity was respiratory rate (N = 2; Barbic et al., 2022; Snow et al., 2019). Finally, those studies that implemented EEG as a data collection technique performed Frequency Band analysis (N = 2; Okamoto et al., 2017; Snow et al., 2019).

From the total of included articles, 5 studies analyzed only one biomarker (Beaudette et al., 2024; Chinazzo et al., 2019; Fanpu et al., 2024; Song et al., 2020; Zhou et al., 2023), while 11 studies incorporated two or more biomarkers (Barbic et al., 2022; Chinazzo et al., 2018; Fischl & Johansson, 2024; Gao et al., 2023; Hu et al., 2022; Okamoto et al., 2017; Roy et al., 2024; Snow et al., 2019; Wang et al., 2018; Wang et al., 2023; Wu & Wagner, 2023).

#### Well-Being Self-Report Measure

The reviewed studies employed a wide range of self-report methods, reflecting the diversity in their objectives and approaches. As shown in Table 5, the most commonly used tools included the Thermal Comfort Vote (TCV), Thermal Sensation Vote (TSV), and other thermal perceptions. The regulations indicated by the American Society of Heating, Refrigerating, and Air-Conditioning Engineers (ASHRAE) were used as guidelines for an important number of articles in this review. In addition to thermal-related measures, some studies incorporated tools to evaluate emotional states, such as the Positive and Negative Affect Schedule (PANAS), or subjective responses like the Stanford Sleepiness Scale and the Self-Assessment Manikin (SAM) for emotional reactions.

Also, it is important to notice that the scales/questionnaires included were not exclusively based on well-being (e.g., psychological well-being). Most of these instruments were primarily focused on assessing participants’ comfort and subjective experience regarding environmental conditions.

## Discussion

The present literature review systematized the current scientific evidence on methodologies used to investigate the impact of the indoor built environment on well-being, focusing on assessing physiological variables (e.g., cerebral and cardiac activity, skin conductance). After completing the systematic searches, 16 articles matched our criteria for inclusion. In the following sections, we will discuss what we consider to be some of the most relevant aspects for advancing research in this field.

### Research field characterization

From the general analysis of the research’s characteristics, it is possible to notice a significant increase in the studies conducted in the field over the past four years (N = 11). This shows a growing interest in exploring and understanding how variables of the indoor built environment can impact physiological and psychological variables in real-life settings. Although this result describes more than 60% of the included studies, it might not be surprising considering that during the COVID-19 pandemic, the limitations regarding lockdown increased the time spent inside the built environment. Hence, the lockdown generated a change in the relationship with the built environment, especially due to the dissolution of the limits and roles of different built environments (i.e., home office, studying from home; (Daniel, 2020; Martin et al., 2022; Xiao et al., 2021). Even when it is not a groundbreaking finding, we believe that this is a feasible proof of how science is grounded in socio-cultural, political, and economic reality (Haggis, 2008; Haraway, 2020). Thus, to elicitate relevant questions for generating useful evidence, science should not be considered or developed in isolation from the context where it takes place (Haraway, 2020). In this sense, in the attempts for science and scientific knowledge to be generalizable, there is a need not to forget the specificity of some context and the need for science to be sensitive enough to consider and embrace diversity.

Science’s contextualization not only refers to the time domain but also to the geographical location. All the included studies were conducted in the Northern Hemisphere, with 50% taking place in China. These results may reflect the increasing concern in these countries regarding indoor environmental quality, particularly issues related to healthy buildings and energy efficiency. Examples include efforts to reduce carbon emissions (Zhou et al., 2023) and decrease energy consumption (Gao et al., 2023). This focus aligns with current standards for healthy buildings (Wang et al., 2023) and supports the pursuit of the United Nations Sustainable Development Goals for 2030 (United Nations, 2024).

### Indoor Environmental Quality Variables

From the results presented, it can be understood that the current models that govern building designs are based on the simplistic assumption that human beings react in a disjointed and monotonous way to the stimuli they are exposed to. To evaluate the comfort and well-being of people in indoor spaces, the research has addressed the assessment of IEQ factors. This has mainly been done separately. Only 3 out of the 16 studies link more than one factor (Barbic et al., 2022; Chinazzo et al., 2019; Wang et al., 2023), and only one study considers three factors (Wang et al., 2023).

None of the eligible articles in this systematic review considered noise as an IEQ variable to study, a finding that stands out given the established role of noise (or unwanted sound) as a risk factor capable of directly or indirectly activating stress pathways, potentially affecting health and well-being (Basner et al., 2014). This omission is notable, as research demonstrates noise exposure’s adverse physiological and psychological effects (Altomonte et al., 2024; Jensen et al., 2018; Medvedev et al., 2015). A possible explanation for this gap may lie in the predominant focus on thermal comfort and energy efficiency observed in the articles included in this review, suggesting a prioritization of these aspects over other critical environmental factors such as acoustic quality. This underscores the need for future studies to adopt a more integrated approach to indoor environmental quality by examining noise alongside thermal comfort, lighting, and air quality. A holistic understanding of well-being requires considering how these factors interact, both within individuals and across different occupants, rather than assessing them in isolation.

The reviewed articles lack a holistic vision of IEQ criteria, focusing mainly on thermal comfort, which appears to be the most influential factor in user performance and behavior (Frontczak & Wargocki, 2011). However, this focus often overlooks the integration of other relevant environmental stimuli. This integration between the senses and their information content is associated with multi-modal phenomena and cross-modal stimuli interaction. The former comprises a combination or union of multiple unisensory and independent inputs, while the term cross-modal refers to situations where a stimulus of one sensory modality is shown to exert an influence on our perception or responsiveness to stimuli presented in another sensory modality (Spence et al., 2009).

The built environment is a complex system characterized by feedback, agent interrelation, and non-linear, discontinuous relationships. However, the empirical results that support its development often focus on objective associations between stimuli and responses without appreciating the complexity of built environments. These contexts involve combinations of continuous and transient exposures and produce multi-layered psychophysiological effects that drive the occupant’s perception and behavior (Bond et al., 2012; Hanc et al., 2019). Determining the cause-and-effect relationships between well-being, comfort, and environmental parameters is complex because the combinations, apart from being multiple, can interact synergistically or cancel each other out antagonistically, influencing the occupants’ physical, physiological and psychological responses (Chu et al., 2004).

Further research is needed that considers the simultaneous interrelation and integration between environmental factors and their effects on people from the technical, social, mental, and physiological construct (Schweiker et al., 2020; Wei et al., 2020). Neither the perception of comfort nor that of well-being in the built environment can be extended linearly from one physical domain to another, and although the evaluation becomes more complex, it must be taken into account that stimuli continuously and interrelatedly influence personal evaluations.

### What does *Well-Being* mean?

The methods used in the reviewed articles to assess self-reported well-being are highly diverse. Most commonly, they involve measures of satisfaction and/or comfort derived from ASHRAE standards, such as the Thermal Sensation Vote (TSV) and Thermal Comfort Vote (TCV). This variability may reflect the lack of consensus on conceptualisations of well-being.

Wellbeing is considered a state that integrates psychological, social, and physical dimensions, influenced by the environment in which individuals find themselves (Altomonte et al., 2024). The term well-being is often understood simplistically as synonymous with wellness, happiness, and quality of life and/or associated with comfort and health (Bluyssen et al., 2011; Ghaffarianhoseini et al., 2018; J. Lee et al., 2011; Rohde et al., 2020). There is currently no consensus on the definition of well-being in the built environment, so there is no straightforward approach to measuring pre-existing or new buildings. However, there are three clear methods for evaluating the indoor and psychosocial environment characteristics that cause psychosocial and physical stress and their influencing factors. These use medical examinations, extensive self-report questionnaires (which are not based exclusively on well-being), and diverse and varied observation and monitoring techniques (Bluyssen et al., 2011), but without the appearance of comparative analysis with neurological variables.

On the other hand, and as mentioned, it is extremely complex to broadly measure well-being due to the complexity of its definition that addresses states related to health, others with perception according to immediate requirements (i.e., comfort), and other subjective and emotional terms such as love or happiness (J. Lee et al., 2011). To respond to the needs and requirements of building users for both temporary comfort and sustained well-being, it is important to advance the definition of parameters and their interactions, including potential compensations between them. This progress will enable us to characterize people’s responses to this multi-modal and cross-modal environmental stimulation and evaluate how these responses and the corresponding behaviors change over time over time, depending on the context and personal characteristics of the individual. Achieving this, however, requires establishing a clear and consensual framework for what should be tracked and measured.

Progress must be made towards a paradigm shift from current methods and metrics that are typically and repeatedly used to evaluate the qualities and performances of a built environment for (or towards) understanding intermodal stimulation and the synergistic integration between different indoor environmental quality parameters from an interdisciplinary perspective.

### Physiology: What should we be looking for?

Considering the diversity of hypotheses, experimental designs, and technical-methodological aspects of the reviewed studies, it is difficult to determine one exclusive biomarker to address the study of the impact of the indoor built environment on well-being. Our results indicate that the type of physiological analysis employed in each study is largely dictated by the chosen data collection technique and the specific research question under investigation. For instance, iButton devices are commonly used for skin temperature measurements, ECG for heart rate and heart rate variability, and EEG for frequency-based analyses of brain activity. However, these physiological signals have largely been treated as isolated measures, with little consideration of their simultaneous interactions within a broader embodied system.

To critically advance the field, we must move beyond fragmented physiological analyses and toward a more integrated understanding of bodily dynamics. Indoor built environment research could benefit from studying physiological activity in the light of brain/body physiological dynamics and interoceptive processes (e.g., using Heartbeat-Evoked Potentials, HEP; Montoya et al., 1993); Movement-Related Potentials (MRP; Hallett, 1994); Blink-Related Potentials (BRP; Wunderlich & Gramann, 2021); Fixation-Related Potentials (FRP; Kaunitz et al., 2014). A clear example is the case of HEP, which has been widely implemented for the study of interoception (Gumilar et al., 2022), a process that has been used for assessing well-being (Craig, 2002; Erle et al., 2021; Pinna & Edwards, 2020; Wheaton et al., 2022). Hence, implementing these types of integrative analyses might be useful in understanding bodily dynamics modulated by specific aspects of the indoor built environment.

However, the inclusion of these complex analyses is not exempt from challenges. To be properly included, research design and analysis changes might be necessary. In this sense, clear hypothesis-driven analytic strategies based on appropriate theoretical frameworks will be required to reduce the dimensionality of potentially unlimited variables (King & Parada, 2021; Parada & Rossi, 2021). An example of how researchers can overcome this challenge is by implementing the *Scalable Experimental Design* (SED) heuristic (Matusz et al., 2019; Parada, 2018; Shamay-Tsoory & Mendelsohn, 2019), which will allow the design and development of classically structured experiments that later can be expanded towards real/virtual environments. One of the main premises of SED is that this type of experimental design will allow parametric testing of reliable neurobehavioral markers (e.g., HEP) in different settings, from laboratory to everyday situations. In short, SED posits the opportunity to study the evolutionary-given sensorimotor possibilities of the encounter between the agent and the environment. This might be relevant for the research field of indoor built environments since it will allow the development of specific research questions regarding the impact of indoor variables -as they naturally occur in the real world- in cognitive (e.g., attention), psychological (e.g., well-being), and even interactional (e.g., environment as a facilitator for social interaction) processes that can be carefully and systematically tested in research. Nevertheless, technical-methodological development is needed not only to move forward in the research field. Even when (neuro) physiological markers or dynamics might be identified and systematically studied, there is still a need for working within a coherent and *unified* theoretical framework to generate connections between empirical data and the existent knowledge and theories (Gramann et al., 2021).

Another important issue regarding physiological analysis, is the necessity of interdisciplinary research teams or collaborations. The importance of having experts from different research fields who can contribute to the analyses and interpretation of the collected data, especially due to the lack of standardized biomarkers for the study field, is increasing. Specifically, including neuroscientists and cognitive scientists will allow us to deal with large and complex databases and move forward to identifying key biomarkers or physiological components to parametrically and systematically incorporate and explore them from classical laboratory settings to the real world.

#### From plan drawing to the real world

Research on the indoor built environment seeks to understand how environmental variables shape diverse aspects of human life: daily activities (e.g., work, study, rest), leisure, and health, among others. Understanding these variables can help us incorporate them into architectural design. In this sense, we identify at least two key elements to consider before conducting research.

First, a need to implement research paradigms that allow the study of everyday dynamics (Parada and Rossi 2024). Designing and incorporating experimental tasks resembling real-world tasks and demands are crucial. From the articles included in this review, the realization of neuropsychological assessments in contexts with different indoor environmental variables being manipulated (e.g., luminosity and air ventilation) were the most common experimental tasks. Even when these are specific and structured tests designed for studying cognitive functioning, it is possible to test how cognitive functioning might be affected by environmental variables since they resemble cognitive load in, for example, everyday work or study-related contexts.

On the other hand, the incorporation of naturalistic research paradigms that emulate real-world activities and neurobehavioral processes with higher degrees of ecological validity (i.e., less restrictive settings than the classical neurocognitive laboratory) can contribute to the understanding of the impact of the environment on cognitive and physiological processes as they naturally take place in the world. Here, the SED approach described before offers the possibility to first develop well-controlled laboratory studies to test experimental factors despite their diminished degree of ecological validity (Parada, 2018). Then, it will provide the chance to conduct real-world experiments with less experimental control but with higher ecological validity regarding action and behavior (Gramann et al., 2021; Parada, 2018). As elegantly outlined by Gramann and collaborators (2017, 2021), the real world turns *into the* laboratory. Regarding our results, some of the included articles implemented naturalistic types of tasks (e.g., book reading and office tasks), which show the intention and motivation for generating insights regarding real-world contexts.

Considering the interest in conducting research with valid and meaningful questions to contribute to the design of the indoor built environment, the development of new technologies for research could not be done at a better time. Almost two decades ago, the Mobile Brain-Body Imaging (MoBI) technical-methodological framework offers the possibility to move the neuroscientific research field from the laboratory to the real world, thanks to the portability of devices and the progress made in terms of managing large and complex data analyses (Jungnickel et al., 2019; Klug & Gramann, 2021; Makeig et al., 2009). MoBI’s portability and advances in real-time data acquisition and processing enable researchers to track cognitive, physiological, and behavioral responses as individuals engage with their surroundings naturally. Hence, By achieving greater theoretical-methodological coherence, the field can move toward integrative, multi-modal research approaches that capture the dynamic, embodied, and situated nature of cognition. Ultimately, this will generate deeper insights into the ways built environments shape cognitive processes, psychological well-being, and health, paving the way for truly evidence-based architectural and environmental design (Parada and Rossi 2024).

## Limitations

This systematic review has some limitations that require consideration for interpreting results.

These limitations stem from the current state of the field and the characteristics of the included studies, rather than the methodological approach of this review itself. First, no eligible articles in the reviewed literature included noise as an Indoor Environmental Quality (IEQ) variable, despite its recognized importance in influencing well-being and comfort in indoor environments. This gap highlights the need for more comprehensive research incorporating noise alongside other IEQ variables to better understand its impact on physiological and psychological states. Second, the studies included in this review were predominantly conducted in the Northern Hemisphere, with no representation from the Southern Hemisphere. This geographical limitation restricts the applicability of findings to regions with different cultural, environmental, and climatic conditions. Third, most studies focused on isolated IEQ factors, such as thermal comfort or lighting, rather than addressing the dynamic interplay between multiple environmental stimuli. This fragmented approach limits the understanding of multimodal and cross-modal effects that are critical for capturing the complexity of human-environment interactions. Lastly, the wide range of self-report methods reported introduces variability in definitions and measurements, complicating the comparability of findings.

## Conclusion

The present systematic review highlights methodological aspects of the indoor built environment research field in combination with neuroscience research. The theoretical-methodological development in neuroscience research (i.e., 3E Cognition/Mobile Brain/Body Imaging framework) can provide insights to generate new research questions, as well as collect, analyze, and interpret larger datasets to comprehend the impact of the indoor build environment on cognition, psychological well-being, and health.

Among the principal results of this review, it is possible to identify a significant relationship between different variables of the indoor built environment and psycho-physiological states, although studies of isolated variables proliferate with little holistic vision that includes multi-modal phenomena and cross-modal stimuli interaction. For instance, thermal comfort was found to be the most commonly studied IEQ variable affecting heart activity and skin temperature. These findings are relevant considering the impact of the environment on the health-disease continuum and everyday life; as the opportunity that they offer to generate strategies for interventions and improving quality of life. Our research also highlights the need to shift toward using advanced technologies, such as Mobile Brain/Body Imaging (MoBI), to capture real-time biometric data in natural settings. These technologies can be extrapolated and applied to everyday scenarios, including health-related, work, and educational contexts.

## Declarations

### Ethics approval and consent to participate

Not applicable.

### Consent for publication

Not applicable.

### Availability of data and materials

The dataset is available at the Open Science Framework (OSF): https://osf.io/qc5tx/

### Competing interests

The authors declare that the present work was conducted in the absence of any potential conflict of interest.

### Funding

AGC, PWM & FJP were supported by the Agencia Nacional de Investigación y Desarrollo de Chile (ANID) through Proyecto de Exploración grant number 13220156. FJP also receives funding by a grant from the Berlin University Alliance 113_MC_GlobalHealth. AGC also receives funding from the Deutsche Forschungsgeneubschaft (DFG, German Research Foundation), project number GRK 2185/2. MAP receives funding from the Agencia Nacional de Investigación y Desarrollo (ANID) 2024-2124108.

### Authors’ contributions

AGC, PW, and FJP conceptualized the present study. AGC designed the study. MAP, AGC, EV, and JMS, carried out the review and data extraction processes. AGC and MAP conducted the analysis of extracted information. AGC wrote the first version of the manuscript. MAP, AGC, PW, and FJP wrote the final version of the manuscript. All authors approved the final version of the manuscript.

## Acknowledgements

The authors would like to thank Rodrigo García Alvarado and Karina Neira Zambrano for their support on this project. Also, the authors would like to thank Valentina Santander and Cristobal Carrasco-Gubernatis for their help during the screening process of the first draft of the present manuscript.

## Footnotes

1 The revision of the 3E Cognition paradigm exceeds the aim of the present review.

2 https://osf.io/qc5tx/

3 This step was conducted without the collaboration of a librarian due to institutional limitations.

4 Q1 Were there clear criteria for inclusion in the case series?; Q2 Was the condition measured in a standard, reliable way for all participants included in the case series?; Q3 Were valid methods used for identification of the condition for all participants included in the case series?; Q4 Did the case series have consecutive inclusion of participants?; Q5 Did the case series have complete inclusion of participants?; Q6 Was there clear reporting of the demographics of the participants in the study?; Q7 Was there clear reporting of clinical information of the participants?; Q8 Were the outcomes or follow up results of cases clearly reported?; Q9 Was there clear reporting of the presenting site(s)/clinic(s) demographic information?; Q10 Was statistical analysis appropriate?

5 Q1 Were the criteria for inclusion in the sample clearly defined?; Q2 Were the study subjects and the setting described in detail?; Q3 Was the exposure measured in a valid and reliable way?; Q4 Were objective, standard criteria used for measurement of the condition?; Q5 Were confounding factors identified?; Q6 Were strategies to deal with confounding factors stated?; Q7 Were the outcomes measured in a valid and reliable way?; Q8 Was appropriate statistical analysis used?

## Notes

https://osf.io/qc5tx/

### Competing Interest Statement

The authors have declared no competing interest.

## References

1. Abdulaali, H. S., Usman, I., Hanafiah, M., Abdulhasan, M., Hamzah, M., & Nazal, A. (2020). Impact of poor indoor environmental quality (IEQ) to inhabitants’ health, wellbeing and satisfaction. International Journal of Advanced Science and Technology, 29(3), 1–14.

2. Alanazy, A. R. M., Wark, S., Fraser, J., & Nagle, A. (2019). Factors Impacting Patient Outcomes Associated with Use of Emergency Medical Services Operating in Urban Versus Rural Areas: A Systematic Review. International Journal of Environmental Research and Public Health, 16(10). 10.3390/ijerph16101728

3. Al horr, Y., Arif, M., Katafygiotou, M., Mazroei, A., Kaushik, A., & Elsarrag, E. (2016). Impact of indoor environmental quality on occupant well-being and comfort: A review of the literature. International Journal of Sustainable Built Environment, 5(1), 1–11.

4. Altomonte, S., Allen, J., Bluyssen, P. M., Brager, G., Heschong, L., Loder, A., Schiavon, S., Veitch, J. A., Wang, L., & Wargocki, P. (2020). Ten questions concerning well-being in the built environment. Building and Environment, 180, 106949.

5. Altomonte, S., Kaçel, S., Martinez, P. W., & Licina, D. (2024). What is NExT? A new conceptual model for comfort, satisfaction, health, and well-being in buildings. Building and Environment, 252(111234), 111234.

6. Ancora, L. A., Blanco-Mora, D. A., Alves, I., Bonifácio, A., Morgado, P., & Miranda, B. (2022). Cities and neuroscience research: A systematic literature review. Frontiers in Psychiatry / Frontiers Research Foundation, 13, 983352.

7. Arksey, H., & O’Malley, L. (2005). Scoping studies: towards a methodological framework. International Journal of Social Research Methodology, 8(1), 19–32.

8. Aromataris, E., & (Eds.)., M. Z. (2020). JBI manual for evidence synthesis. https://scholar.google.com/citations?user=KBsFTz8AAAAJ&hl=en&oi=sra

9. Azzazy, S., Ghaffarianhoseini, A., GhaffarianHoseini, A., Naismith, N., & Doborjeh, Z. (2021). A critical review on the impact of built environment on users’ measured brain activity. Architectural Science Review, 64(4), 319–335.

10. Banaei, M., Ahmadi, A., Gramann, K., & Hatami, J. (2020). Emotional evaluation of architectural interior forms based on personality differences using virtual reality. Frontiers of Architectural Research, 9(1), 138–147.

11. Banaei, M., Hatami, J., Yazdanfar, A., & Gramann, K. (2017). Walking through Architectural Spaces: The Impact of Interior Forms on Human Brain Dynamics. Frontiers in Human Neuroscience, 11. 10.3389/fnhum.2017.00477

12. Barbic, F., Minonzio, M., Cairo, B., Shiffer, D., Cerina, L., Verzeletti, P., Badilini, F., Vaglio, M., Porta, A., Santambrogio, M., Gatti, R., Rigo, S., Bisoglio, A., & Furlan, R. (2022). Effects of a cool classroom microclimate on cardiac autonomic control and cognitive performances in undergraduate students. The Science of the Total Environment, 808, 152005.

13. Bartuska, T. J. (2007). Understanding environment (s): built and natural. The Built Environment: A Collaborative Inquiry into Design and Planning, 33–43.

14. Basner, M., Babisch, W., Davis, A., Brink, M., Clark, C., Janssen, S., & Stansfeld, S. (2014). Auditory and non-auditory effects of noise on health. The Lancet, 383(9925), 1325–1332.

15. Beaudette, E., Foo, E., Islam Molla, M. T., Johnson, K., Dupler, E., Gagliardi, N., Woelfle, H., Halvey, M., & Dunne, L. (2024). Investigating user experience of on-body heating strategies in indoor environments. Ergonomics in Design: The Magazine of Human Factors Applications, 32(2), 25–32.

16. Beemer, C. J., Stearns-Yoder, K. A., Schuldt, S. J., Kinney, K. A., Lowry, C. A., Postolache, T. T., Brenner, L. A., & Hoisington, A. J. (2021). A brief review on the mental health for select elements of the built environment. Indoor and Built Environment, 30(2), 152–165.

17. Beil, K., & Hanes, D. (2013). The influence of urban natural and built environments on physiological and psychological measures of stress--a pilot study. International Journal of Environmental Research and Public Health, 10(4), 1250–1267.

18. Besser, L. M., Rodriguez, D. A., McDonald, N., Kukull, W. A., Fitzpatrick, A. L., Rapp, S. R., & Seeman, T. (2018). Neighborhood built environment and cognition in non-demented older adults: The Multi-Ethnic Study of Atherosclerosis. Social Science & Medicine, 200, 27–35.

19. Beukeboom, C. J., Langeveld, D., & Tanja-Dijkstra, K. (2012). Stress-reducing effects of real and artificial nature in a hospital waiting room. Journal of Alternative and Complementary Medicine, 18(4), 329–333.

20. Bluyssen, P. M., Janssen, S., van den Brink, L. H., & de Kluizenaar, Y. (2011). Assessment of wellbeing in an indoor office environment. Building and Environment, 46(12), 2632–2640.

21. Bodo, T. (2019). Rapid urbanisation: theories, causes, consequences and coping strategies. Annals of Geographical Studies, 2(3), 32–45.

22. Bolouki, A. (2022). Neurobiological effects of urban built and natural environment on mental health: systematic review. Reviews on Environmental Health. 10.1515/reveh-2021-0137

23. Bond, L., Kearns, A., Mason, P., Tannahill, C., Egan, M., & Whitely, E. (2012). Exploring the relationships between housing, neighbourhoods and mental wellbeing for residents of deprived areas. BMC Public Health, 12, 48.

24. Cheng, B., Wunderlich, A., Gramann, K., Lin, E., & Fabrikant, S. I. (2022). The effect of landmark visualization in mobile maps on brain activity during navigation: A virtual reality study. Frontiers in Virtual Reality, 3. 10.3389/frvir.2022.981625

25. Chen, Z., He, Y., & Yu, Y. (2020). Attention restoration during environmental exposure via alpha-theta oscillations and synchronization. Journal of Environmental Psychology, 68, 101406.

26. Chinazzo, G., Wienold, J., & Andersen, M. (2018). Combined effects of daylight transmitted through coloured glazing and indoor temperature on thermal responses and overall comfort. Building and Environment, 144, 583–597.

27. Chinazzo, G., Wienold, J., & Andersen, M. (2019). Daylight affects human thermal perception. Scientific Reports, 9(1), 13690.

28. Chu, A., Thorne, A., & Guite, H. (2004). The impact on mental well-being of the urban and physical environment: an assessment of the evidence. Journal of Public Mental Health, 3(2), 17–32.

29. Clark, A. (2000). Mindware: An introduction to the philosophy of cognitive science. 210. https://psycnet.apa.org/fulltext/2001-14520-000.pdf

30. Codinhoto, R., Tzortzopoulos, P., Kagioglou, M., Aouad, G., & Cooper, R. (2009). The impacts of the built environment on health outcomes. Facilities, 27(3/4), 138–151.

31. Craig, A. D. (2002). How do you feel? Interoception: the sense of the physiological condition of the body. Nature Reviews. Neuroscience, 3(8), 655–666.

32. Daniel, S. J. (2020). Education and the COVID-19 pandemic. Prospects, 49(1-2), 91–96.

33. Day, R. (2008). Local environments and older people’s health: dimensions from a comparative qualitative study in Scotland. Health & Place, 14(2), 299–312.

34. De Jaegher, H., Di Paolo, E., & Gallagher, S. (2010). Can social interaction constitute social cognition? Trends in Cognitive Sciences, 14(10), 441–447.

35. Di Paolo, E., & De Jaegher, H. (2012). The interactive brain hypothesis. Frontiers in Human Neuroscience, 6, 163.

36. Di Paolo, E., Rohde, M., & De Jaegher, H. (2010). Horizons for the enactive mind: Values, social interaction, and play. In Enaction: Towards a new paradigm for cognitive science. books.google.com.

37. Djebbara, Z., Jensen, O. B., Parada, F. J., & Gramann, K. (2022). Neuroscience and architecture: Modulating behavior through sensorimotor responses to the built environment. Neuroscience and Biobehavioral Reviews, 138, 104715.

38. Elliott, S. J., Taylor, S. M., Walter, S., Stieb, D., Frank, J., & Eyles, J. (1993). Modelling psychosocial effects of exposure to solid waste facilities. Social Science & Medicine, 37(6), 791–804.

39. Erle, T. M., Mitschke, V., & Schultchen, D. (2021). Did my heart just leap or sink? The role of personality for the relation between cardiac interoception and well-being. Personality and Individual Differences, 170, 110493.

40. Evans, G. W., Wells, N. M., & Moch, A. (2003). Housing and mental health: A review of the evidence and a methodological and conceptual critique. The Journal of Social Issues, 59(3), 475–500.

41. Fanpu, M., Shou Yi, W., & Hua, F. (2024). Research on the health lighting scheme of university library reading room. Heliyon, 10(19), e38089.

42. Fischl, G., & Johansson, P. (2024). Digital occupancy assessment for lighting evaluation: a pilot study to prepare for real-time research results. Architectural Science Review, 1–10.

43. Frontczak, M., & Wargocki, P. (2011). Literature survey on how different factors influence human comfort in indoor environments. Building and Environment, 46(4), 922–937.

44. Gao, Y., Gao, Y., Shao, Z., & Ren, Y. (2023). The effects of indoor temperature and exercise behavior on thermal comfort in cold region: A field study on Xi’an, China. *Energy (Oxford*, England), 273(127258), 127258.

45. Ghaffarianhoseini, A., AlWaer, H., Omrany, H., Ghaffarianhoseini, A., Alalouch, C., Clements-Croome, D., & Tookey, J. (2018). Sick building syndrome: are we doing enough? Architectural Science Review, 61(3), 99–121.

46. Gramann, K., Fairclough, S. H., Zander, T. O., & Ayaz, H. (2017). Editorial: Trends in Neuroergonomics. Frontiers in Human Neuroscience, 11. 10.3389/fnhum.2017.00165

47. Gramann, K., Ferris, D. P., Gwin, J., & Makeig, S. (2014). Imaging natural cognition in action. International Journal of Psychophysiology: Official Journal of the International Organization of Psychophysiology, 91(1), 22–29.

48. Gramann, K., Gwin, J. T., Ferris, D. P., Oie, K., Jung, T.-P., Lin, C.-T., Liao, L.-D., & Makeig, S. (2011). Cognition in action: imaging brain/body dynamics in mobile humans. Reviews in the Neurosciences, 22(6), 593–608.

49. Gramann, K., McKendrick, R., Baldwin, C., Roy, R. N., Jeunet, C., Mehta, R. K., & Vecchiato, G. (2021). Grand Field Challenges for Cognitive Neuroergonomics in the Coming Decade. Frontiers in Neuroergonomics, 2. 10.3389/fnrgo.2021.643969

50. Gumilar, I., Barde, A., Sasikumar, P., Billinghurst, M., Hayati, A. F., Lee, G., Munarko, Y., Singh, S., & Momin, A. (2022). Inter-brain Synchrony and Eye Gaze Direction During Collaboration in VR. Extended Abstracts of the 2022 CHI Conference on Human Factors in Computing Systems, Article Article 345.

51. Haggis, T. (2008). “Knowledge Must Be Contextual”: Some possible implications of complexity and dynamic systems theories for educational research. Educational Philosophy and Theory, 40(1), 158–176.

52. Hallett, M. (1994). Movement-related cortical potentials. Electromyography and Clinical Neurophysiology, 34(1), 5–13.

53. Hall, S. A., Kaufman, J. S., & Ricketts, T. C. (2006). Defining urban and rural areas in U.S. epidemiologic studies. Journal of Urban Health: Bulletin of the New York Academy of Medicine, 83(2), 162–175.

54. Hanc, M., McAndrew, C., & Ucci, M. (2019). Conceptual approaches to wellbeing in buildings: a scoping review. Building Research and Information, 47(6), 767–783.

55. Haraway, D. (2020). Situated knowledges: The science question in feminism and the privilege of partial perspective. In Feminist theory reader (pp. 303–310). Routledge.

56. Hu, J., He, Y., Hao, X., Li, N., Su, Y., & Qu, H. (2022). Optimal temperature ranges considering gender differences in thermal comfort, work performance, and sick building syndrome: A winter field study in university classrooms. Energy and Buildings, 254, 111554.

57. Hu, M., & Roberts, J. (2020). Built Environment Evaluation in Virtual Reality Environments—A Cognitive Neuroscience Approach. Urban Science, 4(4), 48.

58. Jensen, H. A. R., Rasmussen, B., & Ekholm, O. (2018). Neighbour and traffic noise annoyance: a nationwide study of associated mental health and perceived stress. European Journal of Public Health, 28(6), 1050–1055.

59. Jiang, M., Hassan, A., Chen, Q., & Liu, Y. (2020). Effects of different landscape visual stimuli on psychophysiological responses in Chinese students. Indoor and Built Environment, 29(7), 1006–1016.

60. Jonas, H. (1966). The Phenomenon of Life:: Towards a Philosophical Biology. 303pp. *New York*. Jungnickel, E., Gehrke, L., Klug, M., & Gramann, K. (2019). Chapter 10 - MoBI—Mobile Brain/Body Imaging. In H. Ayaz & F. Dehais (Eds.), *Neuroergonomic**s* (pp. 59–63). Academic Press.

61. Jungnickel, E., & Gramann, K. (2016). Mobile Brain/Body Imaging (MoBI) of Physical Interaction with Dynamically Moving Objects. Frontiers in Human Neuroscience, 10, 306.

62. Karakas, T., & Yildiz, D. (2020). Exploring the influence of the built environment on human experience through a neuroscience approach: A systematic review. Frontiers of Architectural Research, 9(1), 236–247.

63. Kaunitz, L. N., Kamienkowski, J. E., Varatharajah, A., Sigman, M., Quiroga, R. Q., & Ison, M. J. (2014). Looking for a face in the crowd: fixation-related potentials in an eye-movement visual search task. NeuroImage, 89, 297–305.

64. Keis, O., Helbig, H., Streb, J., & Hille, K. (2014). Influence of blue-enriched classroom lighting on students cognitive performance. Trends in Neuroscience and Education, 3(3), 86–92.

65. Kim, J., Yadav, M., Ahn, C. R., & Chaspari, T. (2019, June 13). Saliency detection analysis of pedestrians’ physiological responses to assess adverse built environment features. Computing in Civil Engineering 2019. ASCE International Conference on Computing in Civil Engineering 2019, Atlanta, Georgia. 10.1061/9780784482445.012

66. King, J. L., & Parada, F. J. (2021). Using mobile brain/body imaging to advance research in arts, health, and related therapeutics. The European Journal of Neuroscience. 10.1111/ejn.15313

67. Klepeis, N. E., Nelson, W. C., Ott, W. R., Robinson, J. P., Tsang, A. M., Switzer, P., Behar, J. V., Hern, S. C., & Engelmann, W. H. (2001). The National Human Activity Pattern Survey (NHAPS): a resource for assessing exposure to environmental pollutants. Journal of Exposure Analysis and Environmental Epidemiology, 11(3), 231–252.

68. Klug, M., & Gramann, K. (2021). Identifying key factors for improving ICA-based decomposition of EEG data in mobile and stationary experiments. The European Journal of Neuroscience, 54(12), 8406–8420.

69. Kyselo, M. (2014). The body social: an enactive approach to the self. Frontiers in Psychology, 5, 986.

70. Ladouce, S., Donaldson, D. I., Dudchenko, P. A., & Ietswaart, M. (2016). Understanding Minds in Real-World Environments: Toward a Mobile Cognition Approach. Frontiers in Human Neuroscience, 10, 694.

71. Laland, K. N., Odling-Smee, J., & Feldman, M. W. (2000). Niche construction, biological evolution, and cultural change. The Behavioral and Brain Sciences, 23(1), 131–146; discussion 146–175.

72. Lederbogen, F., Haddad, L., & Meyer-Lindenberg, A. (2013). Urban social stress--risk factor for mental disorders. The case of schizophrenia. Environmental Pollution, 183, 2–6.

73. Lederbogen, F., Kirsch, P., Haddad, L., Streit, F., Tost, H., Schuch, P., Wüst, S., Pruessner, J. C., Rietschel, M., Deuschle, M., & Meyer-Lindenberg, A. (2011). City living and urban upbringing affect neural social stress processing in humans. Nature, 474(7352), 498–501.

74. Lee, J., Je, H., & Byun, J. (2011). Well-Being index of super tall residential buildings in Korea. Building and Environment, 46(5), 1184–1194.

75. Lee, S., Shin, W., & Joo, P. E. (2022). Implications of neuroarchitecture for the experience of the built environment: a scoping review. Archnet-IJAR: International Journal of Architectural Research, 16(2), 225–244.

76. Li, J., Jin, Y., Lu, S., Wu, W., & Wang, P. (2020). Building environment information and human perceptual feedback collected through a combined virtual reality (VR) and electroencephalogram (EEG) method. Energy and Buildings, 224, 110259.

77. Liu, W., Lian, Z., & Liu, Y. (2008). Heart rate variability at different thermal comfort levels. European Journal of Applied Physiology, 103(3), 361–366.

78. Lottrup, L., Grahn, P., & Stigsdotter, U. K. (2013). Workplace greenery and perceived level of stress: Benefits of access to a green outdoor environment at the workplace. Landscape and Urban Planning, 110, 5–11.

79. Makeig, S., Gramann, K., Jung, T.-P., Sejnowski, T. J., & Poizner, H. (2009). Linking brain, mind and behavior. International Journal of Psychophysiology: Official Journal of the International Organization of Psychophysiology, 73(2), 95–100.

80. Mallawaarachchi, B. H., De Silva, M. L., Rameezdeen, R., & Chandrathilaka, S. R. (2012). *Green building concept to facilitating high quality indoor environment for building occupants in Sri Lanka*. http://dl.lib.uom.lk/handle/123/16996

81. Marchand, G. C., Nardi, N. M., Reynolds, D., & Pamoukov, S. (2014). The impact of the classroom built environment on student perceptions and learning. Journal of Environmental Psychology, 40, 187–197.

82. Martin, L., Hauret, L., & Fuhrer, C. (2022). Digitally transformed home office impacts on job satisfaction, job stress and job productivity. COVID-19 findings. *PloS One*, *17*(3), e0265131.

83. Matthews, S. A., & Yang, T.-C. (2010). Exploring the role of the built and social neighborhood environment in moderating stress and health. Annals of Behavioral Medicine: A Publication of the Society of Behavioral Medicine, 39(2), 170–183.

84. Matusz, P. J., Dikker, S., Huth, A. G., & Perrodin, C. (2019). Are We Ready for Real-world Neuroscience? Journal of Cognitive Neuroscience, 31(3), 327–338.

85. Mavros, P., Austwick, M. Z., & Smith, A. H. (2016). Geo-EEG: Towards the Use of EEG in the Study of Urban Behaviour. Applied Spatial Analysis and Policy, 9(2), 191–212.

86. Mavros, P., J Wälti, M., Nazemi, M., Ong, C. H., & Hölscher, C. (2022). A mobile EEG study on the psychophysiological effects of walking and crowding in indoor and outdoor urban environments. Scientific Reports, 12(1), 18476.

87. McGowan, J., Sampson, M., Salzwedel, D. M., Cogo, E., Foerster, V., & Lefebvre, C. (2016). PRESS Peer Review of Electronic Search Strategies: 2015 Guideline Statement. Journal of Clinical Epidemiology, 75, 40–46.

88. McMichael, A. J. (2000). The urban environment and health in a world of increasing globalization: issues for developing countries. *Bulletin of the World Health Organization*. https://www.scielosp.org/article/ssm/content/raw/?resource_ssm_path=/media/assets/bwho/v78n 9/v78n9a07.pdf

89. Medvedev, O., Shepherd, D., & Hautus, M. J. (2015). The restorative potential of soundscapes: A physiological investigation. Applied Acoustics, 96, 20–26.

90. Microsoft Corporation. (2018). *Microsoft Excel* (Version 2019 (16.0)). https://office.microsoft.com/excel

91. Montoya, P., Schandry, R., & Müller, A. (1993). Heartbeat evoked potentials (HEP): topography and influence of cardiac awareness and focus of attention. Electroencephalography and Clinical Neurophysiology, 88(3), 163–172.

92. Moola, S., Munn, Z., Tufanaru, C., Aromataris, E., & Sears, K. (2017). Checklist for case series. Joanna Briggs Inst.

93. Moore, T. H. M., Kesten, J. M., López-López, J. A., Ijaz, S., McAleenan, A., Richards, A., Gray, S., Savović, J., & Audrey, S. (2018). The effects of changes to the built environment on the mental health and well-being of adults: Systematic review. Health & Place, 53, 237–257.

94. Möystad, O. (2017). Cognition and the built environment. Routledge.

95. Mulders, D., de Bodt, C., Lejeune, N., Courtin, A., Liberati, G., Verleysen, M., & Mouraux, A. (2020). Dynamics of the perception and EEG signals triggered by tonic warm and cool stimulation. PloS One, 15(4), e0231698.

96. Munn, Z., Barker, T. H., Moola, S., Tufanaru, C., Stern, C., McArthur, A., Stephenson, M., & Aromataris, E. (2020). Methodological quality of case series studies: an introduction to the JBI critical appraisal tool. JBI Evidence Synthesis, 18(10), 2127–2133.

97. Muthukrishna, M., Bell, A. V., Henrich, J., Curtin, C. M., Gedranovich, A., McInerney, J., & Thue, B. (2020). Beyond Western, Educated, Industrial, Rich, and Democratic (WEIRD) Psychology: Measuring and Mapping Scales of Cultural and Psychological Distance. Psychological Science, 31(6), 678–701.

98. Neale, C., Aspinall, P., Roe, J., Tilley, S., Mavros, P., Cinderby, S., Coyne, R., Thin, N., & Ward Thompson, C. (2020). The impact of walking in different urban environments on brain activity in older people. Cities & Health, 4(1), 94–106.

99. Newen, A., De Bruin, L., & Gallagher, S. (2018). The Oxford Handbook of 4E Cognition. Oxford University Press.

100. Okamoto, T., Tamura, K., Miyamoto, N., Tanaka, S., & Futaeda, T. (2017). Physiological activity in calm thermal indoor environments. Scientific Reports, 7(1), 11519.

101. Parada, F. J. (2018). Understanding Natural Cognition in Everyday Settings: 3 Pressing Challenges. Frontiers in Human Neuroscience, 12, 386.

102. Parada, F. J., & Rossi, A. (2018). If Neuroscience Needs Behavior, What Does Psychology Need? Frontiers in Psychology, 9, 433.

103. Parada, F. J., & Rossi, A. (2021). Perfect timing: Mobile brain/body imaging scaffolds the 4E-cognition research program. The European Journal of Neuroscience, 54(12), 8081–8091.

104. Parada, F., & Rossi, A. (2024). The Potential of Neuroarchitecture and 4E-Cognition: From Microbial Dynamics to Active Environments and Back via Scalable Experimental Designs. ESS Open Archive eprints, 59, 05933745.

105. Pateman, T. (2011). Rural and urban areas: comparing lives using rural/urban classifications. Regional Trends, 43(1), 11–86.

106. Pelgrims, I., Devleesschauwer, B., Guyot, M., Keune, H., Nawrot, T. S., Remmen, R., Saenen, N. D., Trabelsi, S., Thomas, I., Aerts, R., & De Clercq, E. M. (2021). Association between urban environment and mental health in Brussels, Belgium. BMC Public Health, 21(1), 635.

107. Persiani, S. G. L., Kobas, B., Koth, S. C., & Auer, T. (2021). Biometric Data as Real-Time Measure of Physiological Reactions to Environmental Stimuli in the Built Environment. Energies, 14(1), 232.

108. Pinna, T., & Edwards, D. J. (2020). A systematic review of associations between interoception, vagal tone, and emotional regulation: Potential applications for mental health, wellbeing, psychological flexibility, and chronic conditions. Frontiers in Psychology, 11, 1792.

109. Portella, A. A. (2014). Built Environment. In A. C. Michalos (Ed.), *Encyclopedia of Quality of Life and Well-Being Research* (pp. 454–461). Springer Netherlands.

110. Power, M. C., Kioumourtzoglou, M.-A., Hart, J. E., Okereke, O. I., Laden, F., & Weisskopf, M. G. (2015). The relation between past exposure to fine particulate air pollution and prevalent anxiety: observational cohort study. BMJ, 350, h1111.

111. Rhodes, R. E., Saelens, B. E., & Sauvage-Mar, C. (2018). Understanding Physical Activity through Interactions Between the Built Environment and Social Cognition: A Systematic Review. Sports Medicine, 48(8), 1893–1912.

112. Ritchie, H., & Roser, M. (2018). Urbanization. *Our World in Data*. https://ourworldindata.org/urbanization?source=content_type%3Areact%7Cfirst_level_url%3Aarticle%7Csection%3Amain_content%7Cbutton%3Abody_link

113. Rohde, L., Larsen, T. S., Jensen, R. L., & Larsen, O. K. (2020). Framing holistic indoor environment: Definitions of comfort, health and well-being. Indoor and Built Environment, 29(8), 1118–1136.

114. Rojas-Líbano, D., & Parada, F. J. (2019). Body-World Coupling, Sensorimotor Mechanisms, and the Ontogeny of Social Cognition. Frontiers in Psychology, 10, 3005.

115. Rossi, A., Cladera, A. G., Luarte, N., Riillo, A., & Parada, F. J. (2019). El sistema cerebro/cuerpo-en-el-mundo es el objeto de estudio de la ciencia cognitiva en el siglo XXI. *Studies in Psychology*, Estudios de Psicología, 40(2), 377–395.

116. Roy, S., Satvaya, P., & Bhattacharya, S. (2024). Effects of indoor lighting conditions on subjective preferences of task lighting and room aesthetics in an Indian tertiary educational institution. Building and Environment, 249(111119), 111119.

117. Salonen, H., Lahtinen, M., Lappalainen, S., Nevala, N., Knibbs, L. D., Morawska, L., & Reijula, K. (2013). Physical characteristics of the indoor environment that affect health and wellbeing in healthcare facilities: a review. Intelligent Buildings International, 5(1), 3–25.

118. Santos, W. M. D., Secoli, S. R., & Püschel, V. A. de A. (2018). The Joanna Briggs Institute approach for systematic reviews. Revista Latino-Americana de Enfermagem, 26, e3074.

119. Schweiker, M., Ampatzi, E., Andargie, M. S., Andersen, R. K., Azar, E., Barthelmes, V. M., Berger, C., Bourikas, L., Carlucci, S., Chinazzo, G., Edappilly, L. P., Favero, M., Gauthier, S., Jamrozik, A., Kane, M., Mahdavi, A., Piselli, C., Pisello, A. L., Roetzel, A., … Zhang, S. (2020). Review of multi-domain approaches to indoor environmental perception and behaviour. Building and Environment, 176, 106804.

120. Schweizer, C., Edwards, R. D., Bayer-Oglesby, L., Gauderman, W. J., Ilacqua, V., Jantunen, M. J., Lai, H. K., Nieuwenhuijsen, M., & Künzli, N. (2007). Indoor time-microenvironment-activity patterns in seven regions of Europe. Journal of Exposure Science & Environmental Epidemiology, 17(2), 170–181.

121. Shamay-Tsoory, S. G., & Mendelsohn, A. (2019). Real-Life Neuroscience: An Ecological Approach to Brain and Behavior Research. Perspectives on Psychological Science: A Journal of the Association for Psychological Science, 14(5), 841–859.

122. Snow, S., Boyson, A. S., Paas, K. H. W., Gough, H., King, M.-F., Barlow, J., Noakes, C. J., & Schraefel, M. c. (2019). Exploring the physiological, neurophysiological and cognitive performance effects of elevated carbon dioxide concentrations indoors. Building and Environment, 156, 243–252.

123. Song, C., Liu, Y., Zhou, X., Wang, D., Wang, Y., & Liu, J. (2020). Identification of local thermal conditions for sleeping comfort improvement in neutral to cold indoor thermal environments. Journal of Thermal Biology, 87, 102480.

124. Spence, C., Senkowski, D., & Röder, B. (2009). Crossmodal processing. Experimental Brain Research. Experimentelle Hirnforschung. Experimentation Cerebrale, 198(2), 107–111.

125. Steinemann, A., Wargocki, P., & Rismanchi, B. (2017). Ten questions concerning green buildings and indoor air quality. Building and Environment, 112, 351–358.

126. Stephan, A., & Walter, S. (2020). Situated affectivity. In The Routledge handbook of phenomenology of emotion (pp. 299–311). Routledge.

127. Sterelny, K. (2010). Minds: extended or scaffolded? Phenomenology and the Cognitive Sciences, 9(4), 465–481.

128. Stevens, R. G., & Rea, M. S. (2001). Light in the built environment: potential role of circadian disruption in endocrine disruption and breast cancer. Cancer Causes & Control: CCC, 12(3), 279–287.

129. Sullivan, W. C., & Chang, C.-Y. (2011). Mental Health and the Built Environment. In A. L. Dannenberg, H. Frumkin, & R. J. Jackson (Eds.), *Making Healthy Places: Designing and Building for Health, Well-being, and Sustainability* (pp. 106–116). Island Press/Center for Resource Economics.

130. Taylor, S. E., Repetti, R. L., & Seeman, T. (1997). Health psychology: what is an unhealthy environment and how does it get under the skin? Annual Review of Psychology, 48(1), 411–447.

131. Thelen, E., & Smith, L. B. (1995). A dynamic systems approach to the development of cognition and action. Journal of Cognitive Neuroscience, 7(4), 512–514.

132. The World Bank. (2022). *Urban Development*. The World Bank. https://www.worldbank.org/en/topic/urbandevelopment/overview#:~:text=Today%2C%20some%2056%25%20of%20the,people%20will%20live%20in%20cities.

133. Thompson, E. (2010). Mind in Life: Biology, Phenomenology, and the Sciences of Mind. Harvard University Press.

134. Turunen, M., Toyinbo, O., Putus, T., Nevalainen, A., Shaughnessy, R., & Haverinen-Shaughnessy, U. (2014). Indoor environmental quality in school buildings, and the health and wellbeing of students. International Journal of Hygiene and Environmental Health, 217(7), 733–739.

135. Ulrich, R. S. (1981). Natural Versus Urban Scenes: Some Psychophysiological Effects. Environment and Behavior, 13(5), 523–556.

136. United Nations. (2018). *The World’s Cities in* 2018 (No. 1). {United Nations, Department of Economic and Social Affairs Population Dynamics}. https://www.un.org/development/desa/pd/sites/www.un.org.development.desa.pd/files/files/docu ments/2020/Jan/un_2018_worldcities_databooklet.pdf

137. United Nations. (2024). The Sustainable Development Goals. United Nations. https://www.un.org/sustainabledevelopment/

138. van den Berg, A. E., Maas, J., Verheij, R. A., & Groenewegen, P. P. (2010). Green space as a buffer between stressful life events and health. Social Science & Medicine, 70(8), 1203–1210.

139. Varela, F. J., Rosch, E., & Thompson, E. (1991). The Embodied Mind: Cognitive Science and Human Experience. Cambridge, MA: MIT Press.

140. Wang, Y., Wang, Q., Wang, L., Li, F., Weschler, L. B., Huang, J., & Zhang, Y. (2023). Potential benefits of short-term indoor exposure to sweet orange essential oil for relaxation during mental work breaks. Journal of Building Engineering, 78(107602), 107602.

141. Ward Thompson, C., Roe, J., Aspinall, P., Mitchell, R., Clow, A., & Miller, D. (2012). More green space is linked to less stress in deprived communities: Evidence from salivary cortisol patterns. Landscape and Urban Planning, 105(3), 221–229.

142. Weeks, J. R. (2010). Defining Urban Areas. In T. Rashed & C. Jürgens (Eds.), *Remote Sensing of Urban and Suburban Areas* (pp. 33–45). Springer Netherlands.

143. Wei, W., Wargocki, P., Zirngibl, J., Bendžalová, J., & Mandin, C. (2020). Review of parameters used to assess the quality of the indoor environment in Green Building certification schemes for offices and hotels. Energy and Buildings, 209, 109683.

144. Wheaton, M. G., Sütterlin, S., Edwards, D. J., & Wellbeing, P. F. (2022). Background: Interoception and heart rate variability have been found to predict outcomes of mental health and well-being. However, these have usually been investigated independently of one another. Objectives: This systematic review aimed to explore a key gap in the current literature, that being, identifying whether HRV and interoception predict emotional regulation. *Improving Wellbeing in Patients With Chronic Conditions: Theory, Evidence, and Opportunities*. https://books.google.com/books?hl=en&lr=&id=orxpEAAAQBAJ&oi=fnd&pg=PA229&dq=%22interoception%22+AND+%22well-being%22&ots=jcjnKDSoJj&sig=Nv3uvDEzB5cNBkP8NK5UDLWhk0o

145. Wunderlich, A., & Gramann, K. (2021). Eye movement-related brain potentials during assisted navigation in real-world environments. The European Journal of Neuroscience, 54(12), 8336–8354.

146. Wu, Z., & Wagner, A. (2023). Effect of short-term thermal history on thermal comfort and physiological responses: A pilot study. Energy and Buildings, 298(113510), 113510.

147. Xiao, Y., Becerik-Gerber, B., Lucas, G., & Roll, S. C. (2021). Impacts of Working From Home During COVID-19 Pandemic on Physical and Mental Well-Being of Office Workstation Users. Journal of Occupational and Environmental Medicine / American College of Occupational and Environmental Medicine, 63(3), 181–190.

148. Yadav, M., Chaspari, T., Kim, J., & Ahn, C. R. (2018). Capturing and quantifying emotional distress in the built environment. Proceedings of the Workshop on Human-Habitat for Health (H3*): Human-Habitat Multimodal Interaction for Promoting Health and Well-Being in the Internet of Things Era*, Article Article 9.

149. Yao, Y., Lian, Z., Liu, W., Jiang, C., Liu, Y., & Lu, H. (2009). Heart rate variation and electroencephalograph--the potential physiological factors for thermal comfort study. Indoor Air, 19(2), 93–101.

150. Yeo, L. B. (2021). Psychological And Physiological Benefits Of Plants In The Indoor Environment: A Mini And In-Depth Review. International Journal of Built Environment and Sustainability, 8(1), 57–67.

151. Yuen, B., & Nyuk Hien, W. (2005). Resident perceptions and expectations of rooftop gardens in Singapore. Landscape and Urban Planning, 73(4), 263–276.

152. Zainal, N. Z., & Hosni, N. (2022). Effects of Urban Built Environment on Mental Health: A Review. Journal of Cognitive Sciences and Human Development, 8(1), 30–48.

153. Zhou, J., Zhang, X., Xie, J., & Liu, J. (2023). Occupant’s preferred indoor air speed in hot-humid climate and its influence on thermal comfort. Building and Environment, 229(109933), 109933.

154. Zhu, H., Wang, H., Liu, Z., Li, D., Kou, G., & Li, C. (2018). Experimental study on the human thermal comfort based on the heart rate variability (HRV) analysis under different environments. The Science of the Total Environment, *616-617*, 1124–1133.

